# Means, motive, and opportunity for biological invasions: genetic introgression in a fungal pathogen

**DOI:** 10.1101/2021.09.25.461217

**Authors:** Flávia Rogério, Cock Van Oosterhout, Maisa Ciampi-Guillardi, Fernando Henrique Correr, Guilherme Kenichi Hosaka, Sandrine Cros-Arteil, Gabriel Rodrigues Alves Margarido, Nelson S. Massola, Pierre Gladieux

## Abstract

Invasions by fungal plant pathogens pose a significant threat to the health of agriculture ecosystems. Despite limited standing genetic variation, many invasive fungal species can adapt and spread rapidly, resulting in significant losses in crop yields. Here, we report on the population genomics of *Colletotrichum truncatum*, a polyphagous pathogen that can infect more than 460 plant species, and an invasive pathogen on soybean in Brazil. We study the whole-genome sequences of 18 isolates representing 10 fields from two major regions of soybean production. We show that Brazilian *C. truncatum* is subdivided into three phylogenetically distinct lineages that exchange genetic variation through hybridization. Introgression affects 2 to 30% of the nucleotides of genomes and varies widely between the lineages. We find that introgressed regions comprise secreted protein-encoding genes, suggesting possible co-evolutionary targets for selection in those regions. We highlight the inherent vulnerability of genetically uniform crops in the agro-ecological environment, particularly when faced with pathogens that can take full advantage of the opportunities offered by an increasingly globalized world. Finally, we discuss “The Means, Motive, and Opportunity” of fungal pathogens and how they can become invasive species of crops. We call for more population genomic studies because such analyses can help identify geographic areas and pathogens that pose a risk, thereby helping to inform control strategies to better protect crops in the future.

## INTRODUCTION

Understanding the eco-evolutionary and human-associated factors underlying the emergence and spread of fungal plant diseases is essential to the implementation of effective control measures (Hessenauer et al., 2020; Stukenbrock & McDonald, 2008). Population genetics and molecular epidemiology can shed light on both the extrinsic and intrinsic drivers of biological invasions by fungal plant pathogens (Gladieux et al., 2015; Grünwald, McDonald, & Milgroom, 2016). In parallel, molecular ecology and evolution can provide insights into how an invasive species with limited genetic variation can evolve into ecologically successful pest species soon after a founder event.

Fungi can rapidly evolve into devastating invasive species, despite their often low genetic diversity (Ali et al., 2014; De Jonge et al., 2013; Gladieux et al., 2018; Latorre et al., 2020; Stauber, Badet, Prospero, & Croll, 2020). Although this is a characteristic that they share with many other invasive species, some unique aspects of fungal population biology may facilitate rapid evolutionary changes and enhance their invasive potential. Despite regular demographic bottlenecks (e.g., during winters), fungal populations generally have periods of huge census size, which can substantial evolutionary changes (Gladieux et al., 2015).

Fungal reproduction is complex and can include both sexual and asexual stages (Alexopoulos, Mims, & Blackwell, 1996). Asexual reproduction can be accomplished through mitotic spores (conidia), which augments the propagule pressure during invasion in non-native habitats. In addition, fungi produce hyphae which are thread-like (filamentous) structures, and this gives fungi a second mode of asexual reproduction, i.e., through mycelial fragmentation. Complementing these asexual modes, during sexual reproduction, compatible individuals may exchange genetic information during plasmogamy and karyogamy between gametes (Alexopoulos et al., 1996). Meiotic spores promote the creation of novel genotypes through recombination, and they serve as dispersal and survival structures (Taylor, Jacobson, & Fisher, 1999). If compatible gametes are derived from genetically diverged lineages, the resulting genetic exchange can lead to genetic introgression.

The large census size of invasive fungal populations enables rapid adaptations to new varieties of resistant plants or antifungals molecules (Barton, 2010; Gladieux et al., 2015). This may be particularly important for fungal populations found on major widespread crops, which due to their vast population sizes, benefit from a high input of novel variation by mutations. These new pathogen genotypes can rapidly spread through genetically uniform host populations. In other words, the high evolvability of some fungi during biological invasions is not necessarily realized through their high standing genetic variation at the point of entry (cf. Fisher’s Fundamental Theorem (Price, 1972)). Rather, many fungi are highly potent biological invaders due to input of *de novo* allelic and genotypic variation every generation, which is a consequence of their high potential for gene flow, recombination and mutation. These drivers of genetic variation play a crucial role in the co-evolutionary arms race between fungal pathogens and their hosts. A sudden increase in these drivers can shift the co-evolutionary balance, and these effects can be particularly severe in an eco-agriculture setting with genetically relatively uniform host plants and animals (Van Oosterhout, 2021).

Multiple populations of the same fungal species can coexist on one host whilst competing for limited resources (Bueno-Sancho et al., 2017; Fournier, Gladieux, & Giraud, 2013; Hartmann, Mcdonald, & Croll, 2018; Hubbard et al., 2015; Persoons et al., 2017; Silva, Várzea, Paulo, & Batista, 2018; Stauber et al., 2020; Thierry et al., 2020; Vieira, Silva, Várzea, Paulo, & Batista, 2018). In the absence of temporal, spatial or habitat barriers, coexistence on the same host may foster genetic exchanges between fungal lineages. If coexisting populations represent previously geographically isolated lineages that have not evolved strong pre- or postzygotic barriers, such introgression can rapidly generate novel genotypic variation. In turn, this can increase the amount of phenotype variation – a phenomenon known as transgressive segregation – which provides more novel substrate for natural selection (Nichols et al., 2015).

Admixture between multiple coexisting populations can also lead to a so-called bridgehead effect, in which highly adapted lineages emerge through recombination among propagules established in an area of first introduction (Bertelsmeier & Keller, 2018; Dutech et al., 2012; Stauber et al., 2020). Ongoing fungal invasions offer a unique opportunity to learn about the ecology and evolution of biotic interactions in human-altered ecosystems (Gladieux et al., 2014; Parker & Gilbert, 2018; Thrall, Hochberg, Burdon, & Bever, 2007; Thrall et al., 2011), and such studies are important to assess the risks posed by pathogens to crop in agriculture.

Anthracnose, mainly associated with the fungus *Colletotrichum truncatum* (Hyde et al., 2009), is one of the most prominent foliar diseases of soybean. This ascomycete is seed-transmitted and can infect more than 460 plant species, including important crops in the Fabaceae and Solanaceae families (Cannon, Damm, Johnston, & Weir, 2012; Damm, Woudenberg, Cannon, & Crous, 2009; Weidemann, TeBeest, & Cartwright, 1988). In Brazil, the worldwide leader in soybean production, previous population genetic studies showed that *C. truncatum* is a recently introduced invasive species structured into three highly divergent clusters coexisting in soybean fields. This suggests there have been multiple introductions from distinct source populations, which are yet to be identified (Rogério, Gladieux, Massola, & Ciampi-Guillardi, 2019).

Here, we use whole-genome resequencing and a population genomics approach to characterize the genetic makeup and infer the evolutionary history of *C. truncatum* causing soybean anthracnose in Brazil. We document evidence of extensive introgression between the three lineages that have invaded Brazilian soybean. Our study highlights the risk that Brazilian *C. truncatum* may represent as a bridgehead for future invasions of soybean-producing areas, facilitating admixture between the three lineages (as well as with any unsampled lineages and possible future immigrant lineages). We discuss why fungi have the means to become potent biological invaders of crops, arguing this is due to their ability to rapidly generate novel genetic variation, in combination with their high propagule pressure accomplished through two models of asexual reproduction. We furthermore discuss why the large biomass of relative uniform crops provides the motive, and the bridgehead populations that enable genetic introgression the opportunity for such biological invasions.

## MATERIALS AND METHODS

### Fungal isolates, DNA extraction and genome sequencing

We used 18 isolates of *Colletotrichum truncatum* from naturally infected commercial soybean fields from Mato Grosso (MT) and Goiás (GO) states in Brazil. Isolates were randomly selected from the three genetic clusters (C1, C2, and C3) previously identified based on the population genetics analysis of microsatellite variation (Rogério, Gladieux, Massola, & Ciampi-Guillardi, 2019) (Table 1). Fungal genomic DNA was extracted using Wizard Genomic DNA Purification kit (Promega) from fresh mycelium grown on potato dextrose liquid medium (Difco). Paired-end libraries were prepared and sequenced on Illumina HiSeq2000 (2×150 bp, insert size ∼ 500 pb) by Genewiz (South Plainfield, USA). The raw reads were deposited at the NCBI/Genbank under the Sequence Read Archive (SRA) accession numbers SAMN13196067 to SAMN13196084 (Table S1).

**Table 1.** *Colletotrichum truncatum* isolates used in this study

| Cluster | Isolate | State | Geographical coordinates | Cultivar | Sampling year |
| --- | --- | --- | --- | --- | --- |
| C1 | LFN0169 | Mato Grosso | 13°18946.799S-56°02933.499W | Y-70 Pioner | 2016 |
| C1 | LFN0185 | Mato Grosso | 13°24939.499S- 56°04904.099W | 7709 Nidera | 2017 |
| C1 | LFN0262 | Mato Grosso | 11°58958.599S- 55°30928.699W | 7709 Nidera | 2017 |
| C1 | LFN0309 | Goiás | 17°27936.299S- 51°07910.399W | 7739 Monsoy | 2017 |
| C1 | LFN0360 | Goiás | 17°45951.599S- 51°02906.999W | PP7200Macro | 2017 |
| C1 | LFN0297 | Goiás | 17°24943.199S- 50°57929.099W | Nidera 5909 | 2017 |
| C1 | LFN0346 | Goiás | 17°45951.599S- 51°02906.999W | Nidera 5909 | 2017 |
| C2 | LFN0205 | Mato Grosso | 13°24939.499S- 56°04904.099W | 8372 Nidera | 2017 |
| C2 | LFN0217 | Mato Grosso | 11°55922.899S- 55°37900.099W | 7709 Nidera | 2017 |
| C2 | LFN0248 | Mato Grosso | 11°58958.599S- 55°30928.699W | 7709 Nidera | 2017 |
| C2 | LFN0318 | Goiás | 17°27936.299S- 51°07910.399W | PP7200Macro | 2017 |
| C2 | LFN0349 | Goiás | 17°45951.599S- 51°02906.999W | Nidera 5909 | 2017 |
| C2 | LFN0288 | Goiás | 17°24943.199S- 50°57929.099W | 7739 Monsoy | 2017 |
| C3 | LFN0150 | Mato Grosso | 13°10910.199S- 56°04908.099W | 8766 Monsoy | 2017 |
| C3 | LFN0225 | Mato Grosso | 11°55922.899S- 55°37900.099W | 8372 Nidera | 2017 |
| C3 | LFN0268 | Mato Grosso | 11°58958.599S- 55°30928.699W | 7709 Nidera | 2017 |
| C3 | LFN0291 | Goiás | 17°24943.199S- 50°57929.099W | PP7200Macro | 2017 |
| C3 | LFN0308 | Goiás | 17°27936.299S- 51°07910.399W | Nidera 5909 | 2017 |

### Read mapping and SNP calling

Read quality was checked using Fastqc (https://www.bioinformatics.babraham.ac.uk/projects/fastqc/). Raw Illumina reads were trimmed for adapter contamination and bases with an average Phred score smaller than 30 were removed using Cutadapt V1.16 software (M. Martin, 2011). Reads were mapped against the reference genome of the previously assembled isolate CMES1059 (Rogério et al., 2020) using BWA-mem v0.7.15 (options -n=5) (Li & Durbin, 2009). Alignments were sorted with Samtools v1.3 (Li et al., 2009), and reads with mapping quality below 30 were removed. Duplicates were removed using Picard v2.7 (http://broadinstitute.github.io/picard/). Single nucleotide polymorphisms (SNPs) and indels were called using the HaplotypeCaller module from the Genome Analysis Toolkit v4.0.12 (GATK) (McKenna et al., 2010), with the option *–emitRefConfidence* GVCF. The gVCF files listing variants were merged using CombineGVCFs and genotyped using GenotypeGVCFs. Monomorphic sites were included using the argument *include_nonvariantsites*. High confidence SNPs were identified using GATK’s VariantFiltration module, following GATK’s best practices (http://www.broadinstitute.org/gatk/guide/best_practices), with parameters: QD < 2.0 (Variant Quality), FS > 60.0 (Phred score Fisher’s test), MQ < 40.0 (Mapping Quality), MQRankSum < 12.5 (Mapping Quality of Reference reads vs alternative reads) and ReadPosRankSum < 8.0 (Distance of alternative read from the end of the reads).

### Population structure

We used Splitstree v4 (Huson & Bryant, 2006) to visualize relationships between isolates in a phylogenetic network based on pseudoassembled genomic sequences generated from the tables of SNPs with the reference sequence as a template. We also used the pairwise homoplasy index (PHI) test implemented in Splitstree to test the null hypothesis of clonality; recombination is expected to result in exchangeable sites within lineages.

Population structure was analyzed using methods optimized for the analysis of large datasets and which do not assume Hardy-Weinberg equilibrium. We performed principal components (PCA) and discriminant analysis of principal components (DAPC) in R with the package Adegenet v2.0 (Jombart & Ahmed, 2011), using the Find.Clusters function. DAPC is a non-model-based method using PCA as a prior step, which provides a description of clusters using discriminant functions. We retained the first 20 principal components. DAPC identifies an optimal number of genetic clusters that best describes the data by running a k-means clustering algorithm and comparing the different clustering solutions using the Bayesian Information Criterion (BIC). Population structure was also analyzed using the software SNMF (Frichot, Mathieu, Trouillon, Bouchard, & François, 2014) which estimates ancestry coefficients based on sparse non-negative matrix factorization and least squares optimization. We calculated ancestry coefficients for 2 to 10 ancestral populations (*K*) using 100 replicates for each *K*. The preferred number of *K* was chosen using a cross-entropy criterion based on the prediction of masked haplotypes to evaluate the error of ancestry estimation. All clustering analyses were based on biallelic SNPs without missing data.

### Diversity and divergence

Polymorphism and divergence statistics were computed in 100 kb non-overlapping windows using the Scikit-allel V1.0.2 Python package (Miles & Harding, 2017). Summary statistics were plotted for the 10 the largest contig using Circos v0.67 software (Connors et al., 2009).

### Recombination analyses

Recombination events were analyzed using the software RDP4 (Martin, Murrell, Golden, Khoosal, & Muhire, 2015) implementing seven independent detection algorithms: RDP (Martin & Rybicki, 2000), Geneconv (Padidam, Sawyer, & Fauquet, 1999), Bootscan (Martin, Posada, Crandall, & Williamson, 2005), MaxChi (Smith, 1992) Chimaera (Posada & Crandall, 2002), SiScan (Gibbs, Armstrong, & Gibbs, 2000) and 3Seq (Boni, Posada, & Feldman, 2007). Whole sequences of the ten largest contigs were scanned using the default settings for the window size. Tests were conducted using a critical value α = 0.05 and p-values were Bonferroni corrected for multiple comparisons of sequences. The evidence for a recombination signal was considered to be strong if it was found to be significant with three or more detection methods. Only events for which the software identified the parental sequences (i.e., no ‘unknowns’) without ambiguous start and end position of the recombination block were considered.

We used PopLddecay version v3.4 (Zhang, Dong, Xu, He, & Yang, 2019) to investigate the patterns of linkage disequilibrium decay within *C. truncatum* genetic groups as coefficient of linkage disequilibrium (*r*^2^) (Hill & Robertson, 1968) calculated for all pairs of SNPs less than 300 kb apart. For this, we used biallelic SNPs, excluding missing data and sites with minor allele frequencies below 10%.

### Genome scan for signature of genetic exchanges

We used Chromopainter v0.0.4 (Lawson, Hellenthal, Myers, & Falush, 2012) for probabilistic chromosome painting to infer recent shared ancestry between *C. truncatum* lineages. This method “paints” individuals in “recipient” populations as a combination of segments from “donor” populations, using linkage disequilibrium information for probability computation and assuming that linked alleles are more likely to be exchanged together during recombination events. We ran three separate analyses, each considering one particular *C. truncatum* lineage as a collection of haplotypes to be painted, and all lineages as donors. The recombination scaling constant *N*_*e*_ and emission probabilities (μ) were calculated as averages weighted by contigs’ length determined by LDhat (Auton & McVean, 2007). Estimates of these parameters for each lineage were obtained by running the expectation maximization algorithm with 200 iterations. These analyses were based on biallelic SNPs dataset without missing data.

Fine-scale admixture between *C. truncatum* lineages also was analyzed using the software HYBridCheck (Ward & van Oosterhout, 2016), which uses a sliding window to scan for sudden changes in nucleotide divergence between sequences, thus identifying potential genetic exchanges where nucleotide divergence is significantly lower. The similarities were visualized through a plot employing the primary colors red, green, and blue, using the 100 bp windows based on the proportion of SNPs shared between the pairwise sequences, with a stepwise increment of 1 bp. In cases where all SNPs are shared between just two of the three lineages, the hybrid color is an exact 50% mix of two primary colors. Hence, yellow, purple, and turquoise colors pinpoint regions of possible recent genetic exchange between two sequences. We carried out this analysis on a triplet involving one isolate representative of each lineage: isolates LFN0297 (lineage C1), LFN0318 (lineage C2), and LFN0308 (lineage C3) for the ten largest contigs.

HybridCheck and Chromopainter identify genomic regions of shared ancestry, and such signal can be caused either by genetic introgression, or by incomplete lineage sorting (Durand, Patterson, Reich, & Slatkin, 2011). In order to differentiate between genetic introgression and incomplete lineage sorting, we also used HybridCheck to estimate the age of recombinant regions. If the genomic region coalescence before the split of the species (or lineages), the signal is consistent with incomplete lineage sorting. However, if the coalescence event is dated after the speciation event (or after the bifurcation of the lineages in the tree), the genetic exchange has occurred after the divergence. In the latter case, the signal is consistent with genetic introgression after hybridization (Jouet, McMullan, & Van Oosterhout, 2015). Recombination blocks were then dated assuming a strict molecular clock with a mutation rate of 10^−8^ per generation, assuming a generation time of one year.

### Functional enrichment

To characterize genes present in the introgressed regions between *C. truncatum* lineages, we extracted the transcripts and proteins from the corresponding regions in the reference genome (Rogério et al., 2020) using GffRead (Pertea & Pertea, 2020). We used SignalP v5.0 (Armenteros et al., 2019) to identify secreted proteins. GO terms were assigned from re-annotated transcripts using Blast2go (Conesa et al., 2005) against the NCBI non-redundant database and used in the enrichment analysis.

### Demographic inferences

To infer the evolutionary history of the genetic lineages we used the Python package Dadi (Gutenkunst, Hernandez, Williamson, & Bustamante, 2009). The method implemented in Dadi infers demographic parameters based on a diffusion approximation to the site frequency spectrum (SFS). The Python script easySFS.py (available at https://github.com/isaacovercast/easySFS) was used to convert the VCF file into a three-dimensional joint site frequency spectrum (3D-JSFS). The SFS was folded because no appropriate outgroup was available. We compared twelve demographic models including strict isolation, isolation with migration (asymmetrical migration rates), and isolation with population size changes, with four possible topologies, using the demographic modeling workflow (*dadi_pipeline*) from Portik et al. 2017 (Fig. S1). For each model, we performed four rounds of optimizations; for each round, we ran multiple replicates and used parameter estimates from the best scoring replicate (highest log-likelihood) to seed searches in the following round. We used the default settings in *dadi_pipeline* for each round (replicates = 10, 20, 30, 40; maxiter = 3, 5, 10, 15; fold = 3, 2, 2, 1), and optimized parameters using the Nelder-Mead method (*optimize_log_fmin*). We used the optimized parameter sets of each replicate to simulate the 3D-JSFS, and the multinomial approach was used to estimate the log-likelihood of the given the model. We assessed the model’s goodness-of-fit by maximizing the model likelihood and visual inspection of the residuals between the site frequency spectra generated by the inferred model and the real data.

## RESULTS

### Population structure and levels of genetic variation

Read mapping and variant calling for 18 isolates of *C. truncatum* submitted to whole-genome sequencing identified 2,220,191 biallelic Single Nucleotide Polymorphisms (SNPs), distributed across 128 contigs (see Table S1 to sequencing statistics). To assess population subdivision and to visualize relationships among isolates we built a neighbor-net network with Splitstree based on the full set of SNPs. This phylogenetic network revealed three groups, henceforth referred to as “lineages” C1, C2 and C3 (Fig. 1A). Lineage C1 was connected to the rest of the dataset by a long, non-reticulated branch consistent with relatively long-term genetic isolation. Lineage 2 and 3 were connected by branches showing extensive reticulations (looping in the network) indicating a history of recombination or incomplete lineage sorting (Fig. 1A). In a Discriminant Analysis of Principal Component (DAPC) modelling *K*=2 to *K*=10 populations, the Bayesian Information Criterion monotonously decreased with increasing *K*, preventing clear choice of a best supported model, but the composition of clusters identified at *K*=3 matched what was observed in the neighbor-net network (Fig. 1B and Fig. S2). Likewise clustering by sparse non-negative matrix factorization algorithms, as implemented in the SNMF method, identified *K* = 3 as the best supported model based on cross-entropy. The ancestry coefficients estimated with sNMF revealed essentially the same pattern of population subdivision as the DAPC and neighbor-net network. However, three isolates (LFN0318, LFN0217, and LFN0150) shared ancestry in two clusters, suggesting admixture between lineages (Fig.1C and Fig.S2).

**Figure 1.**
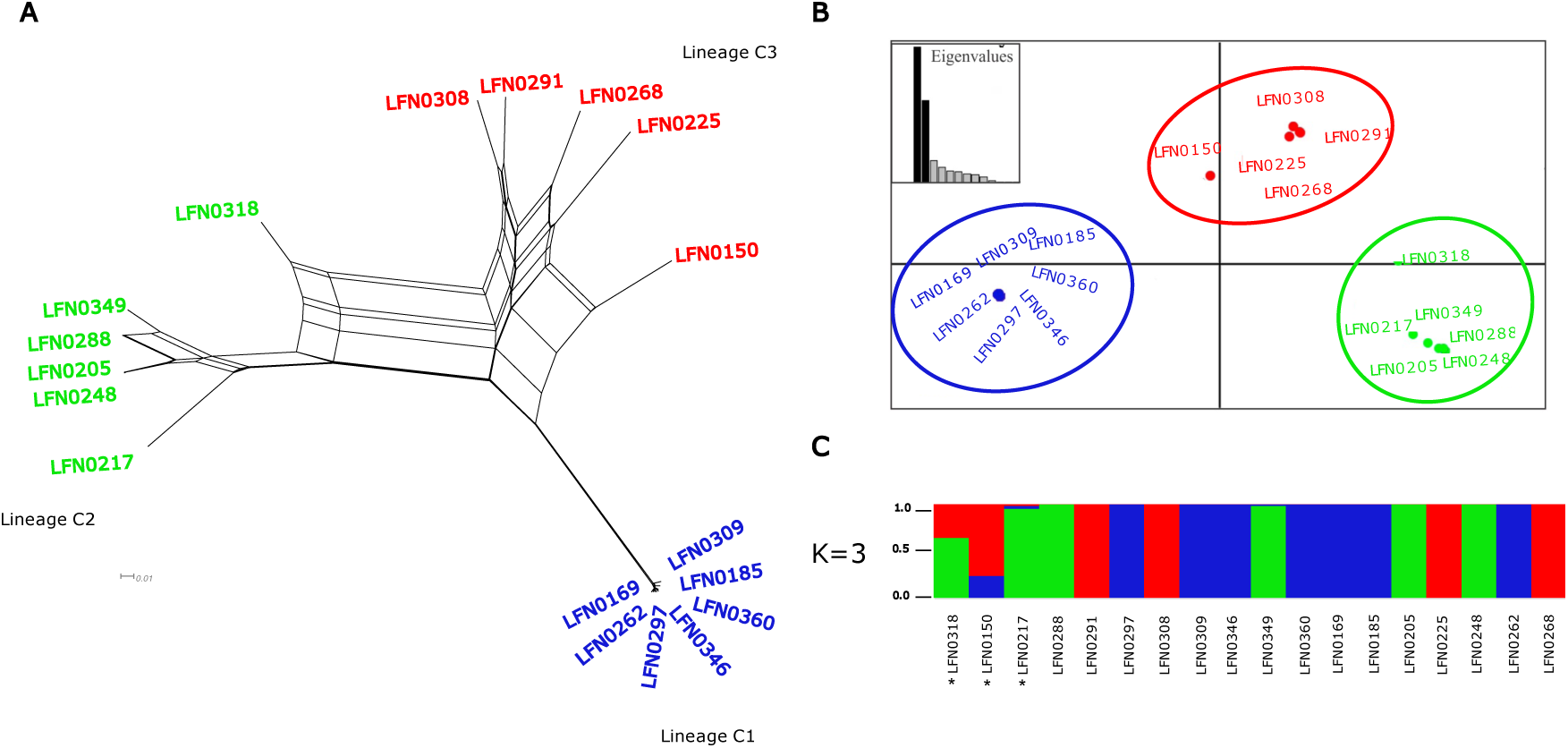
Population subdivision of *Colletotrichum truncatum*. (A) Neighbor-Net networks showing relationships between isolates identified on the basis of the full set of SNPs without missing data. The groups revealed are referred to as lineage C1, C2, and C3. (B) Scatterplot from discriminant analysis of principal components (DAPC). (C) Individual ancestry coefficients estimated using sNMF. Each isolate is represented by a thick vertical line in the most probably number of groups (K=3), and bar colors represent each lineage. Asterisks indicate admixture isolates.

Nucleotide diversity was nearly twice as high in C3 than in C2, and it was more than one order of magnitude higher in C2 than in C1 (C3: π=0.0113/bp; C2: π=0.0062/bp; C1: π=0.0002/bp; Table 2; Fig.2). In lineage C2, regions of relatively high nucleotide diversity were interspersed with tracts of low diversity (Fig. S3). Tajima’s D values were either close to zero, or they were negative in the three lineages (C1: D=0.008; C2: D=-0.380; C3: D=-0.177; Table 2); a negative value is consistent with population expansion after a recent bottleneck or founder event. Absolute divergence (*dxy*) among lineages was similar between the three pairs of lineages (dxy=0.018/bp between C1 and C2, 0.018/bp between C2 and C3, and 0.015/bp between C1 and C3; Fig. 2).

**Table 2.** Summary of genomic diversity within *Colletotrichum truncatum* lineages in nonoverlapping 100kb windows

| Lineage | N<sup>a</sup> | π<sup>b</sup> | D<sup>c</sup> |
| --- | --- | --- | --- |
| C1 | 7 | 0.0002 (0.0003) * | 0.008 (1.006) |
| C2 | 6 | 0.0062 (0.0041) | -0.380 (0.858) |
| C3 | 5 | 0.0113 (0.0056) | -1.177 (0.528) |

<sup>a</sup> Sample size
<sup>b</sup> Nucleotide diversity per base pair, for all contigs larger than 100kb
<sup>c</sup> Tajima's neutrality statistic, for all contigs larger than 100kb
\* Standard deviation

**Figure 2.**
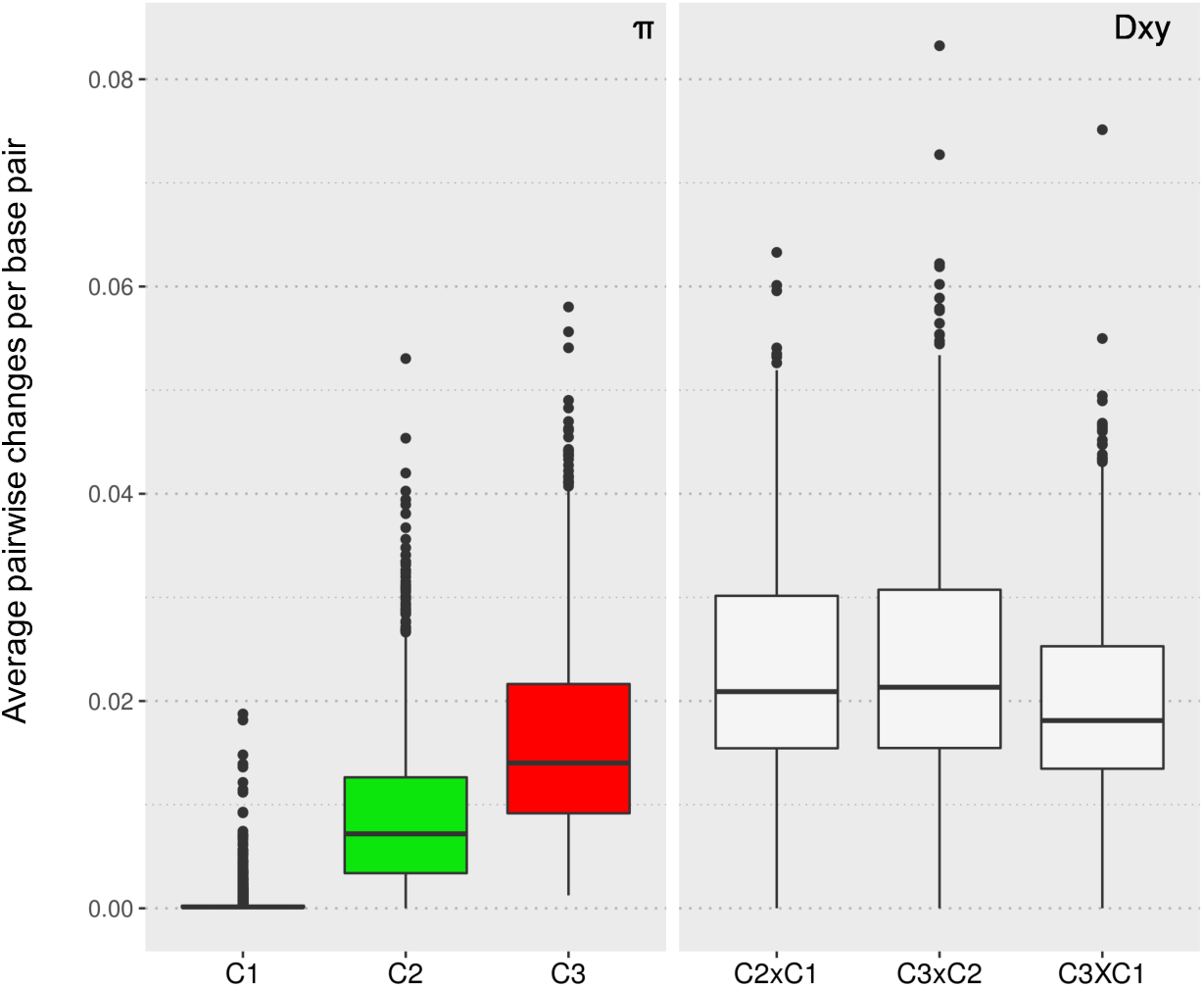
Box plots of the average populations pairwise nucleotide changes per site in 100-kb windows within (nucleotide diversity (*π*)) and between lineages (*dxy*).

### Footprints of recombination

Recombination was analyzed in the ten largest contigs of each lineage, covering ∼30% of the reference assembly. We detected a total of 375 recombined blocks using RDP4 software on the contigs analyzed, with stretches of nucleotide similarity across lineages distributed in a block-like structure (Table S2). Recombination rates differed significantly across lineages; based on the total of 375 recombination events, lineage C3 was found to have received the highest number of recombination events (n=203), followed by C2 (n=148), and with C1 receiving significantly fewer events (n=24) (Randomization test: p<10^−6^; Table S2). In this analysis, we counted the number of cases in which C1, C2 or C3 was the recombinant in Table S2. Analyses of linkage disequilibrium showed that LD decayed to half of its maximum value in less than 1kb in lineages C2 and C3, while LD decay was markedly slower and more jagged in C1 (Fig. 3). The large number of recombination events would have homogenized the nucleotide diversity and broken up any LD blocks in C2 and C3, resulting in a smooth LD decay. In contrast, the LD decay is more erratic in C1 because the few recombination events have not managed to break-up all LD blocks. Finally, the PHI test rejected the null hypothesis of clonality in all three lineages (*P* < 0.001).

**Figure 3.**
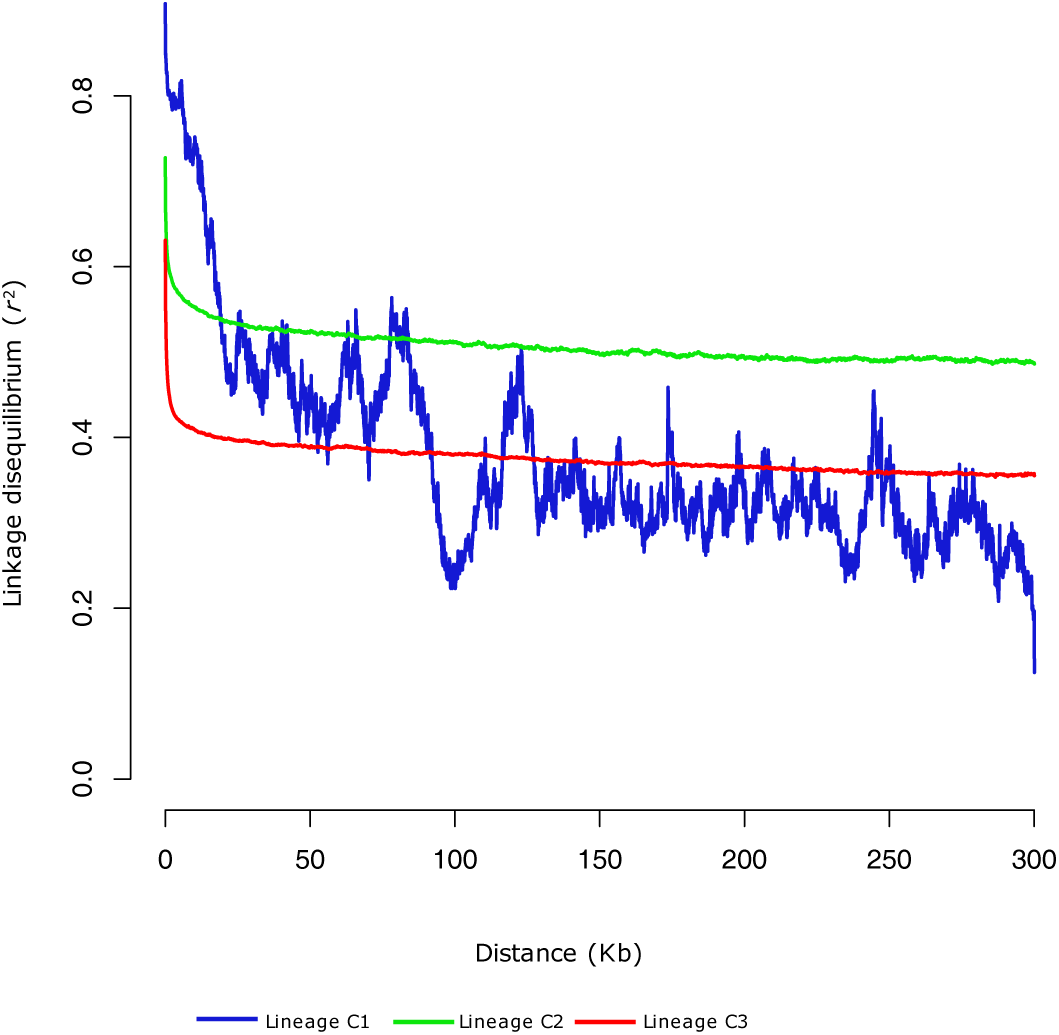
Linkage disequilibrium (LD) decay plots of three genetic lineages of *Colletotrichum truncatum* (C1, C2, and C3). LD measured was calculated for all pairs of SNPs less than 300 kb distance apart.

### Genome scan for signatures of genetic exchanges

Probabilistic chromosome painting revealed genomic regions of shared ancestry between lineages, with shared fragments of size longer than the longest contig in the reference genome (2.37 Mb) (Fig. S4). Regions of shared ancestry were not strictly restricted to the three isolates (LFN0318, LFN0217, and LFN0150) previously detected by sNMF (Fig.1). For lineages C1 and C2, the majority of mutations were assigned to self (i.e., to their cluster of origin), but in some contigs relatively large regions were assigned to other lineages. In lineage C3, mutations tended to have non-zero membership probabilities in multiple clusters. This implies that these polymorphisms are shared across multiple lineages, probably reflecting the extremely high recombination rate shown by this lineage. However, also in lineage C3, some regions were clearly assigned to lineage C1 and self. For contig66, the isolate LFN0318 – a representative of lineage C2 – shared high genetic similarity with lineage C3, consistent with many recent genetic exchanges between these lineages.

Further analyses using HybridCheck revealed a mosaic-like genome structure with well-defined blocks of high nucleotide similarity (Fig. S5). For contig66, relatively few short blocks of high similarity were detected between lineage C1 (using LFN0297 as the representative isolate) and the two other lineages (spanning from 1.4 to 1.6 Mb), while large blocks were detected between lineage C2 and C3 (using LFN0318 and LFN0308 as the representative isolates, respectively) (Fig. 4). Assuming that the contigs analyzed are representative of the rest of the genome, the proportion of genome introgression between lineages varied markedly: 2.4% between C1 (LFN0297) and C2 (LFN0318); 12.7% between C1 (LFN0297) and C3 (LFN0308); and 28.7% between C2 (LFN0318), and C3 (LFN0308) (Table 3 and Table S3).

**Figure 4.**
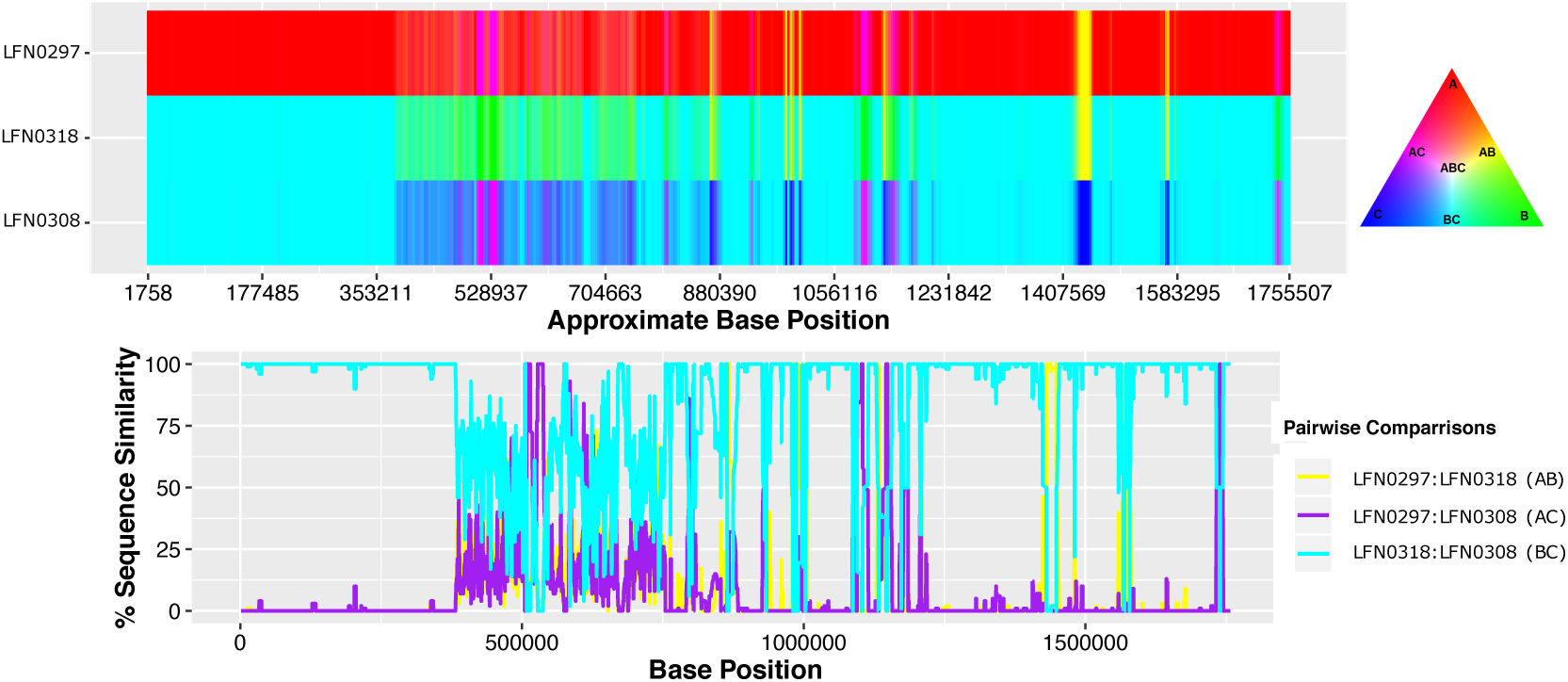
Scan for signature of genetic exchanges between *C. truncatum* lineages. The isolates LFN0297 (lineage C1), LFN0318 (lineage C2), and LFN0308 (lineage C3) were used as representative of each lineage. The sequence similarity along contig 66 among isolates visualized through RBG color triangular in the software HybridCheck. Areas where two sequences have the same color (yellow, purple or turquoise) are indicative of two lineages sharing the same polymorphisms. The bottom panel shows the linear plot of the proportion of SNPs shared between the three pairwise comparisons.

**Table 3.**
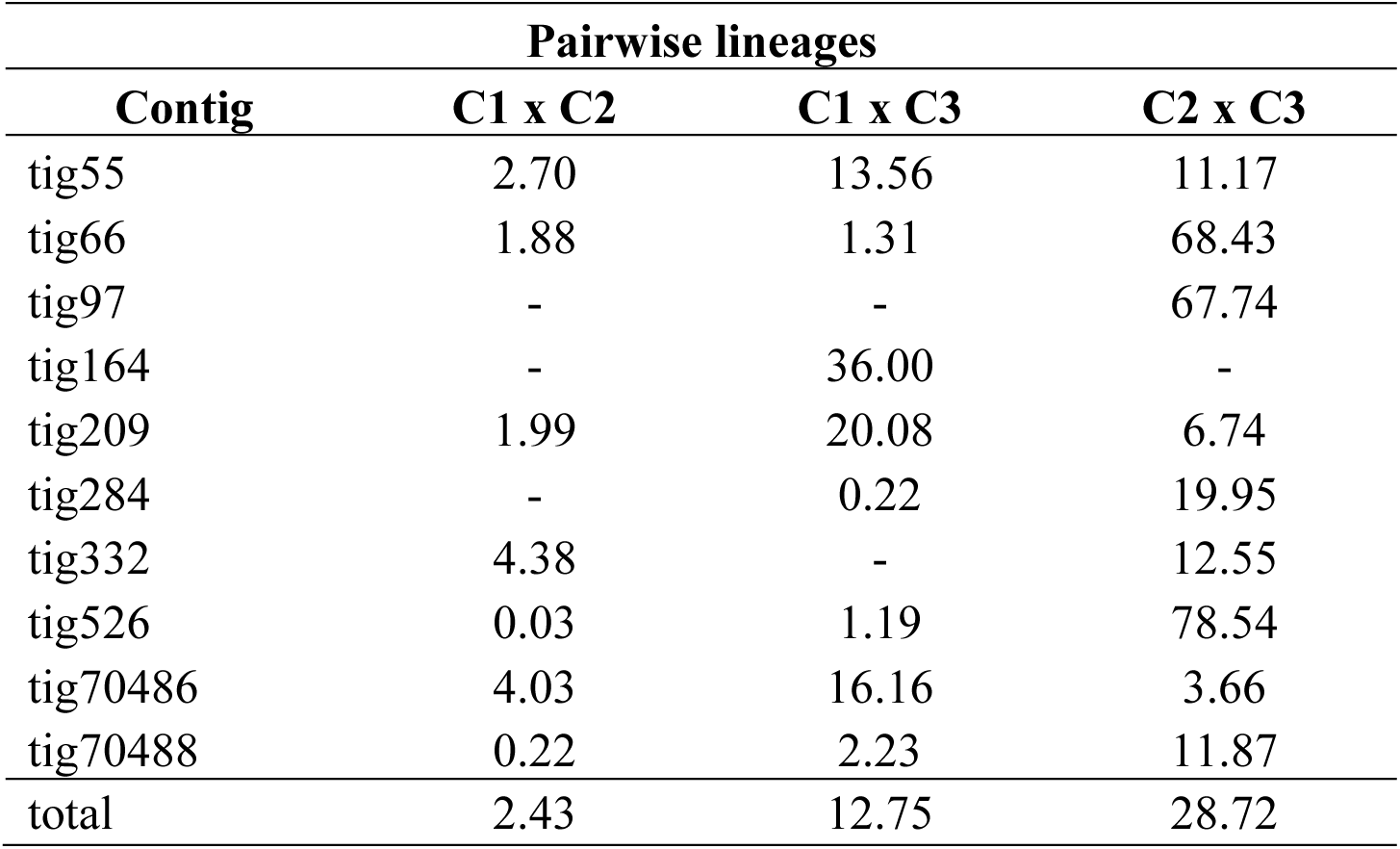
Percentage of genetic introgression between lineages of *Colletotrichum truncatum*

In order to discriminate between incomplete lineage sorting and hybridization after secondary contact, we dated regions of high nucleotide identity between *C. truncatum* lineages detected by HybridCheck. The age estimates of recombinant blocks along the 10 largest contigs revealed recent introgression events. The most recent hybridization event was dated back to 6,100 years before present, assuming a generation time of one year (Table S4). Variation in the age of introgression events was however extensive, most likely because of a lack of closest-related donors for all recombinant regions detected (see Jouet, McMullan, & Van Oosterhout, 2015). The most recent events were more likely to reflect ongoing genetic exchanges. The older events are more consistent with incomplete lineage sorting, or alternatively, the coalescence time may not accurately reflect the timing of genetic introgression because the “true” donors have not been sampled.

In the introgression regions, we identified 357 genes between lineage C1 (LFN0297) and lineage C2 (LFN0318), 389 genes between lineage C1 (LFN0297) and lineage C3 (LFN0308), and 584 genes between lineage C2 (LFN0318) and C3 (LFN0308). The introgressed regions include many secreted protein-encoding genes (between 38 and 62) (Table S5), including proteases and hydrolases which are known virulence-associated factors in pathogens (Monod et al., 2002; Soanes, Richards, & J.Talbot, 2007). However, these regions are not significantly enriched for those genes (binomial test p>0.05), and hence, we must conclude that genetic exchanges between these lineages are not more likely to involve genomic regions with virulence genes. Gene Ontology (GO) analysis revealed enrichment of GO terms between lineages, and these results are reported in Table S6.

### Demographic inferences

To infer the demographic history of the three genetic lineages of *C. truncatum*, we compared three scenarios of isolation with or without migration for the four possible branching orders among lineages, using a diffusion approximation to the SFS implemented using Dadi (We compared 3 × 4 = 12 models compared in total). Note that in the context of the Dadi analysis, the term “migration” is similar to “genetic introgression” in the recombination analysis. Likelihood ratio tests indicated that the model with trifurcating lineages (topology 1) and asymmetrical migration was the most supported (Fig. S6). These results corroborate our recombination analyses. To convert demographic parameter estimates to physical units, we estimated the ancestral population size (N_AB1_) (Fig.5). We used the population mutation rate θ=4N_AB1_μL, where μ was assumed to be approximately 1e-8 per generation (Lynch, 2010) and L was the genome size (∼55.1 Mb). This ancestral population size was then used to transform time estimates from Dadi (in units of 2N_AB1_) into calendar years. Divergence was estimated to have initiated about 960,000 years ago, considering a generation time of a year. The lowest population size was estimated for lineage C1, consistent with a severe bottleneck (nu1=2085 individuals), and/or the least input of genetic variation though recombination or genetic introgression. The migration rate was one order of magnitude higher between lineages C2 and C3 (m23=2.25e-6, m32=1.80e-7), than between lineages C1 and C3 (m13=1.05e-6, m31=0.43e-6), and between lineages C1 and C2 (m12=0.49e-6, m21=0.37e-6). Furthermore, the rate of migration into C1 (m21 and m31) was lower than in all other directions. These demographic analyses thus support the recombination and introgression analyses reported above.

**Figure 5.**
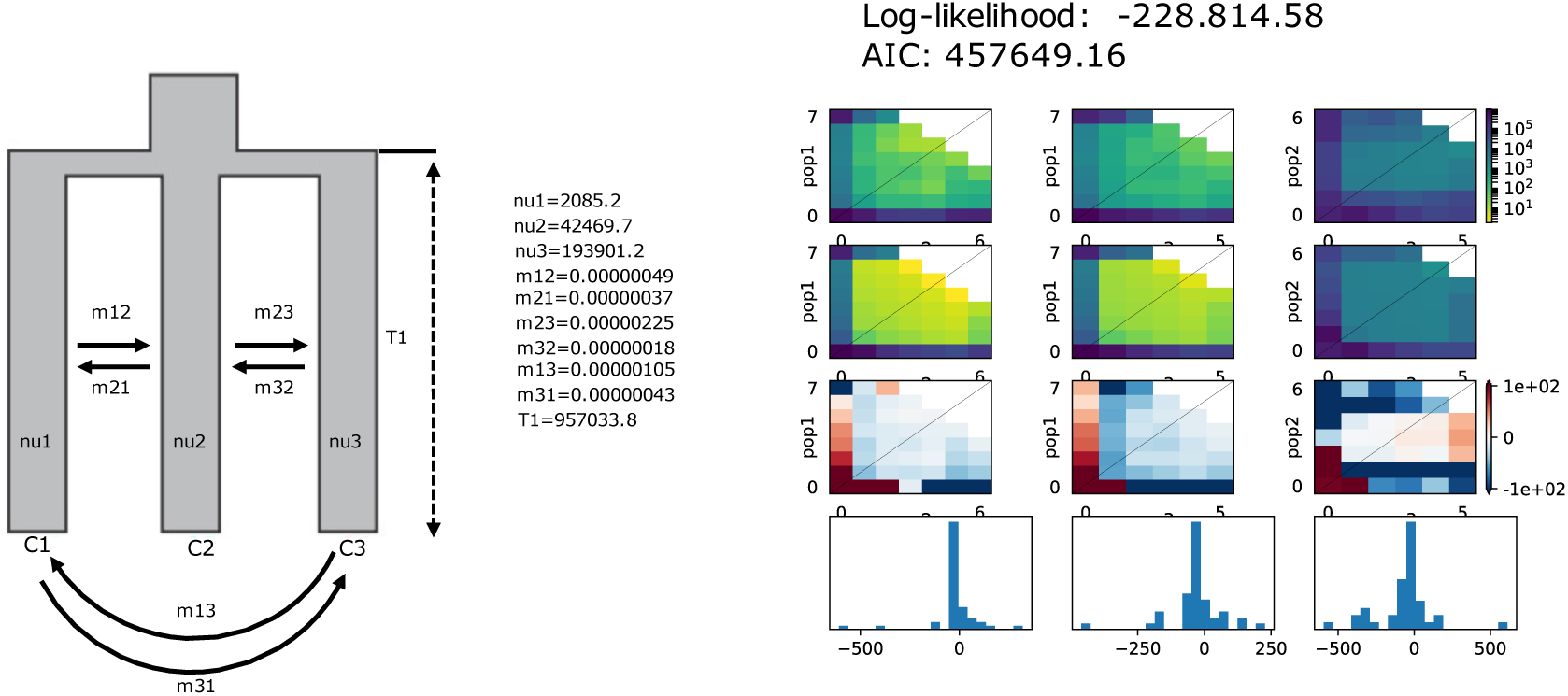
The best support demographic model (isolation with asymmetrical migration) using the three-dimensional site frequency spectrum (3D-SFS) between *Colletotrichum truncatum* lineages (C1, C2, and C3). The following parameters were estimated: <u>nu1</u>: size of C1 after split; <u>nu2</u>: size of C2 after split; <u>nu3</u>: size of C3 after split; <u>m12</u>: asymmetrical migration between C1 and C2; <u>m21</u>: asymmetrical migration between C2 and C1; <u>m23</u>: asymmetrical migration between C2 and C3; <u>m32</u>: asymmetrical migration between C3 and C2.; <u>m13</u>: asymmetrical migration between C1 and C3; <u>m31</u>: asymmetrical migration between C3 and C1; <u>T1</u>: scaled time between the split and the size change (in units of 2*Na generations).

## Discussion

The fungus *Colletotrichum truncatum* is an invasive pathogen on soybean crops in Brazil that causes severe yield losses. We used a population genomics approach to characterize the genetic makeup and infer the evolutionary history of *C. truncatum* using isolates representing two important regions of soybean production in Brazil. We showed that Brazilian *C. truncatum* is subdivided into three phylogenetically equidistant lineages. These lineages possess markedly different levels of standing genomic variation, which could reflect differences in the magnitude of bottlenecks associated with introduction events. A non-exclusive alternative hypothesis is that such differences have been caused by variation in the levels of recombination (i.e., genetic exchanges within a lineage) and/or genetic introgression (genetic exchanges between lineages). All our recombination analyses supported that the lineage C3 with the highest nucleotide diversity was most affected by such genetic exchanges, and that a larger number of these events also affected more genes. Conversely, C1 with the lowest nucleotide variation and the highest level of linkage disequilibrium was least affected by genetic exchanges. Furthermore, these conclusions are corroborated by our demographic analysis which showed that migration (or gene flow) into C1 (m21 and m31) is lower than into all other directions, and that migration into C3 was the highest. Next, we will discuss the evolutionary genomics of *C. truncatum*, the significance of the Brazilian bridgehead population, and the potential of fungal pathogens to evolve into invasive species.

### Evolutionary genomics of C. truncatum

Our clustering analyses supported the existence of three lineages, in agreement with the pattern of population subdivision previously detected based on multilocus microsatellite typing (Rogério, Gladieux, Massola, & Ciampi-Guillardi, 2019). These phylogenetically equidistant lineages are characterized by markedly different levels of genetic standing variation. Genome-wide analyses of variability showed that lineage C1 is largely clonal, reproductively more isolated, and genetically depauperate. Lineage C1 was almost free from introgression, whilst between ∼10 to 30% of assembled contigs of lineage C2 and C3 comprised recombinant (or introgressed) regions. The low level of genetic variation in C1 is consistent with the low number of recombinant blocks (only 6.4%), but it can also indicate a more recent introduction into the country. It means that C1 lineage may simply not have had the opportunity to engage in many genetic exchanges yet. Such recent invasions could be associated with contaminated or infected soybean seeds imported from the U.S. during the 1960s and 1970s (Arantes and Miranda 1993; Hirimoto and Vello 1986; Wysmierski and Vello 2013). Regions of low nucleotide diversity in this lineage corresponded with negative Tajima’s D values, which is consistent with rapid population expansion after a recent founder event. These observations lend further support to our demographic inference of a recent invader, the limited amount of introgression, and the lower effective population size (*Ne*) of C1 lineage.

By contrast, lineage C3 showed a significantly higher level of genetic diversity, which may have been generated over time by genetic recombination and its much larger *Ne* (Note that the large *Ne* estimate may simply be a consequence of the ample genetic variation that has introgressed into this lineage). This lineage may have already been present in Brazil, prior to the introduction of soybean. It is possible that this lineage may have been infecting other host species, such as lima bean and weeds, as proposed by earlier studies (Rogério, Gladieux, Massola, & Ciampi-Guillardi, 2019; Tiffany & Gilman, 1954). If C3 was the first lineage that established itself, it would also have had more opportunity for recombination and genetic introgression than both other lineages, which could have augmented its genetic diversity. Such monopoly effect may also have given this lineage a head-start, both in the co-evolutionary arms race with its host (Van Oosterhout, 2021), as well in the competition with other lineages.

We found evidence of a history of recombination both within and between lineages, applying a combination of integrated approaches. Fine-scale admixture mapping revealed that introgression occurred between the *C. truncatum* lineages coexisting in sympatry despite their relatively deep genomic divergence. The inference of individual ancestry coefficients using probabilistic chromosome painting detected large genomic regions of shared ancestry among the genetic lineages, suggesting relatively recent hybridization. History of recombination and genetic introgression was also supported by the analyzes with HybridCheck, which detected introgressed blocks that differed markedly in age. By estimating the coalescence time of introgressed regions, we found some events have occurred as recently as 6,100 years ago. This suggests that hybridization between *C. truncatum* lineages is a relatively recent - if not ongoing - process. In this analysis, we used a sexual generation time of a year, which is typical in plant pathogenic fungi from temperate areas. However, considering the climate conditions of Brazil, the generation time may be much shorter than one year, and hence, we may have overestimated the age of introgression events. Furthermore, it is unlikely we sampled the actual parental sequence, which would cause a further overestimation of the age. (When identifying the wrong parental sequence, the SNPs that differentiate the parent and the recipient sequence are assumed to have accumulated since recombination took place, erroneously placing the recombination event further in the past). In other words, hybridization events and genetic exchanges may be considerably more recent than our estimate. This is a potentially systematic bias typical for recombination studies, and this can be corrected for by broader (or more intense) sampling.

Although our analyses identified recent introgression between lineages, a substantial proportion of the shared ancestry observed between lineages appear to be caused by incomplete lineage sorting (or alternatively, we overestimated the age). In such cases, the recombination blocks pre-date the lineage divergence (Durand et al., 2011; McMullan et al., 2015), which implies that they are shared ancestral polymorphisms, or that the genetic exchanges occurred before the split of the lineages. *Colletotrichum truncatum* genomes therefore appear to be mosaics of distinct gene genealogies with markedly varied coalescence times. Alternatively, we may not have captured all extant lineages, which would have overestimated the introgression events. Future studies with a more comprehensive sampling may be able to shed further light on this. Next, we discuss our finding in the context of fungi as invasive species.

### Evolutionary genomics of fungi as invasive species

Fungal reproductive biology is conductive for genetic exchange, and such recombination events could both results from sexual reproduction or parasexual events via hyphal anastomosis. The latter mode has already been described for other *Colletotrichum* species (Roca, Davide, Mendes-Costa, & Wheals, 2003; Rosada et al., 2010; Souza-Paccola, Fávaro, Casela, & Paccola-Meirelles, 2003; Vaillancourt, Wang, Hanau, Rollins, & Du, 2000). Given that such genetic exchanges were found in all genomes – i.e., no pure genomes were found – introgression is likely to have augmented both the genetic diversity and the fitness of these hybrid genotypes. Therefore, we would conclude that adaptive introgression may have enhanced the evolutionary potential of *C. truncatum* during its invasion. However, when we tested this hypothesis, we did not find significant enrichment of secreted protein-encoding genes in the introgressed regions. In hindsight, this may not be surprising; in a coevolutionary arms race, genetic novelty at single virulence gene introduced by recombination could provide a selective advantage that helps the recombinant lineage to establish itself (Van Oosterhout, 2021). Indeed, specific targets are likely to be under positive selection, rather than the total number of introgressed genes (Aguileta, Refrégier, Yockteng, Fournier, & Giraud, 2009). In other words, our study may not have discovered “the smoking gun”, but we have established “the means”, i.e., the large number of secreted protein-encoding genes that are exchanged during genetic introgression, which are possible co-evolutionary targets for selection. This implies that introgression can provide the genetic variation required in a host-parasite arms race.

The level of phylogenetic divergence among lineages, coupled with the demographic modeling carried out in this study, enables us to infer that the *C. truncatum* lineages significantly diversified before their joint introduction into Brazil. Based on the ancient signature of some of the recombination events, it is possible that genetic exchanges have occurred during the divergence process. These genetic exchanges would have prevented the accumulation of intrinsic postzygotic barriers, which underpin reproductive isolation in many recently diverged species (Bomblies et al., 2007; Lee et al., 2008; Masly & Presgraves, 2007). Another possibility is that they initially evolved in allopatry, but their later introduction in the same areas may have provided opportunities for secondary contact and hybridization, preventing the accumulation of postzygotic barriers. In summary, the absence of reproductive isolation may have provided ample evolutionary opportunities after secondary contact, allowing for hybridization between diverged lineages.

Introduced populations can overcome consequences of low genetic variation from founders, for instance, through the purging of deleterious alleles during bottlenecks events, and via the fixation of *de novo* beneficial mutations from standing variation (Estoup et al., 2018; Frankham, 2005; Schrieber & Lachmuth, 2017). The genetic and environmental homogeneity found in soybean fields is thought to favor a huge census size of invasive fungal populations. Furthermore, without genetic diversity in the host, pathogens can rapidly spread and fix *de novo* evolved adaptations, overcoming new resistant varieties of crops and fungicides applications. Hybridization could be particularly important in this context because admixture could promote adaptation by rapidly creating novel allelic combinations (Hessenauer et al., 2020; McMullan et al., 2015; Nader et al., 2019). We propose that admixture has elevated the amount of genotypic variation generated by genetic introgression. In turn, this could have increased the amount of phenotypic variation due to transgressive segregation, providing novel substrate for natural selection (Nichols et al., 2015).

Bridgehead population may enhance the adapted invasive potential of species by enabling genetic exchanges between diverged lineages in the areas of first introduction. In this scenario, bridgehead populations may acquire new traits increasing the probability of successful establishment and further spread relative to native population (Bertelsmeier & Keller, 2018). We hypothesize that the admixture between *C. truncatum* lineages may lead to a bridgehead population, producing a highly adapted invasive population. Although the bridgehead effect has been proposed as a potential explanation for many successful biological invasions (Gau, Merz, Falloon, & Brunner, 2013; Leduc et al., 2015; van Boheemen et al., 2017) there is currently no clear empirical support for this hypothesis (but see Simon et al., 2011 for an example of an invasive fish species). To contain biological invasions, vigilance for invasive bridgehead populations is needed since they have the potential to generate new introductions (Bertelsmeier & Keller, 2018) and increase the adaptive evolutionary potential through genetic reassortment during hybridization.

Our study reinforces the practical applications of population genomics in preventing or curtailing, pathogen dissemination by supporting early interventions to limit economic damage (Stam et al., 2021). The Brazilian *C. truncatum* may represent a risk as a bridgehead for future invasions of soybean-producing areas. Our data highlights the inherent vulnerability of genetically uniform crops in the agro-ecological environment, particularly when faced with pathogens that can take full advantage of the opportunities offered by an increasingly globalized word. Some fungi have “The Means, Motive and Opportunity” to become invasive pathogens of crops. Many fungal pathogens possess the means in the form of a high propagule pressure through two modes of asexual reproduction, as well as the ability to rapidly generate novel genotypic variation, particularly through genetic introgression. Some fungi also have a motive, given the large biomass of genetically near-uniform crops that are their natural host plants. Few species also have the opportunity in the form of bridgehead populations that enable the genetic exchange that fuel the co-evolutionary arms race. Invasive crop pathogens, like *C. truncatum*, have “The Means, Motive and Opportunity” to pose the greatest risk to future food security. Population genomics can help identify pathogens that pose such risk, thereby helping to inform control strategies to better protect crops in the future.

## Acknowledgements

The authors are grateful for the financial support given by São Paulo Research Foundation (FAPESP, Grand/Award Number:2017/09178-8), National Science and Technology Development Council (CNPq, Grand/Award Number: 153958/2016-2), and National Council for the Improvement of Higher Education (CAPES/PDSE, Grand/Award Number: 88881.133223/2016-01, PROEX/CAPES, Grand/Award Number: 330002037002P3).

## Data accessibility

DNA sequences: Short Read Archive Accession in Table S1 (Supporting information)

## Author Contributions

M.C.G., and N.S.M.J conceived and designed the research; F.R., and M.C.G collected the samples; S.C.A. obtained genomic data; F.H.C., G.K.H., and G.R.A.M. performed genetic analysis; F.R., C.V.O., and P.G. analyzed the data and wrote the manuscript.

