## Supplementary figures and images for "Means, motive, and opportunity for biological invasions: genetic introgression in a fungal pathogen"

### Figure S1

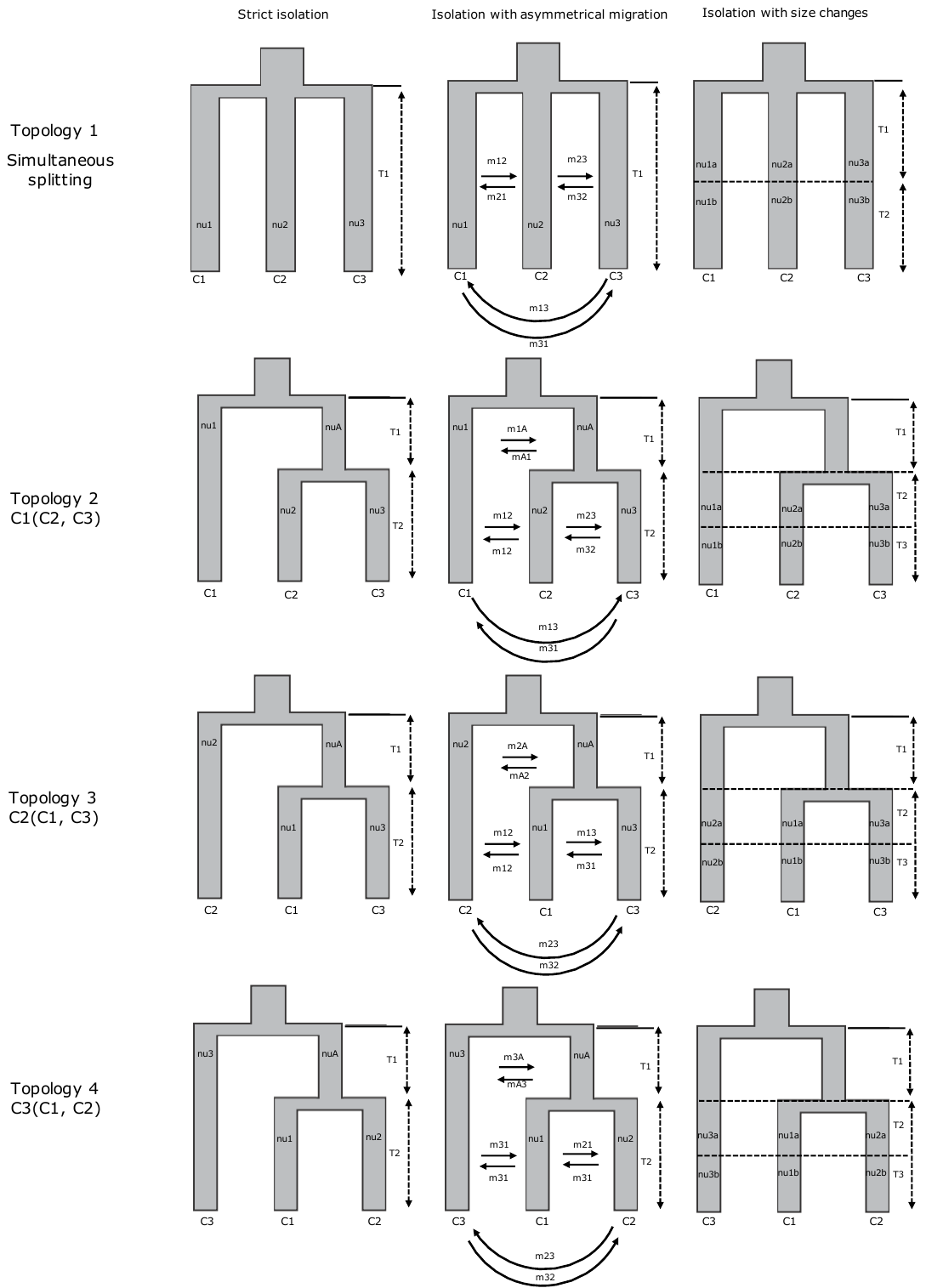

**Figure S1.** Visual representation of the 3D demographic models fitted.
