## Supplementary material for "Means, motive, and opportunity for biological invasions: genetic introgression in a fungal pathogen": Figure S2

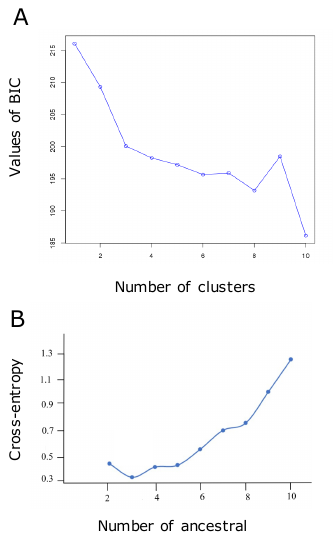

**Figure S2.** (A) Bayesian information criteria (BIC) indicating the most probable number of genetic groups by discriminant analysis of principal components analysis (DAPC). (B) Cross-entropy as a function of the number of clusters K modeled in sNMF analyses of population subdivision.
