## Supplementary material for "Means, motive, and opportunity for biological invasions: genetic introgression in a fungal pathogen": Figure S3

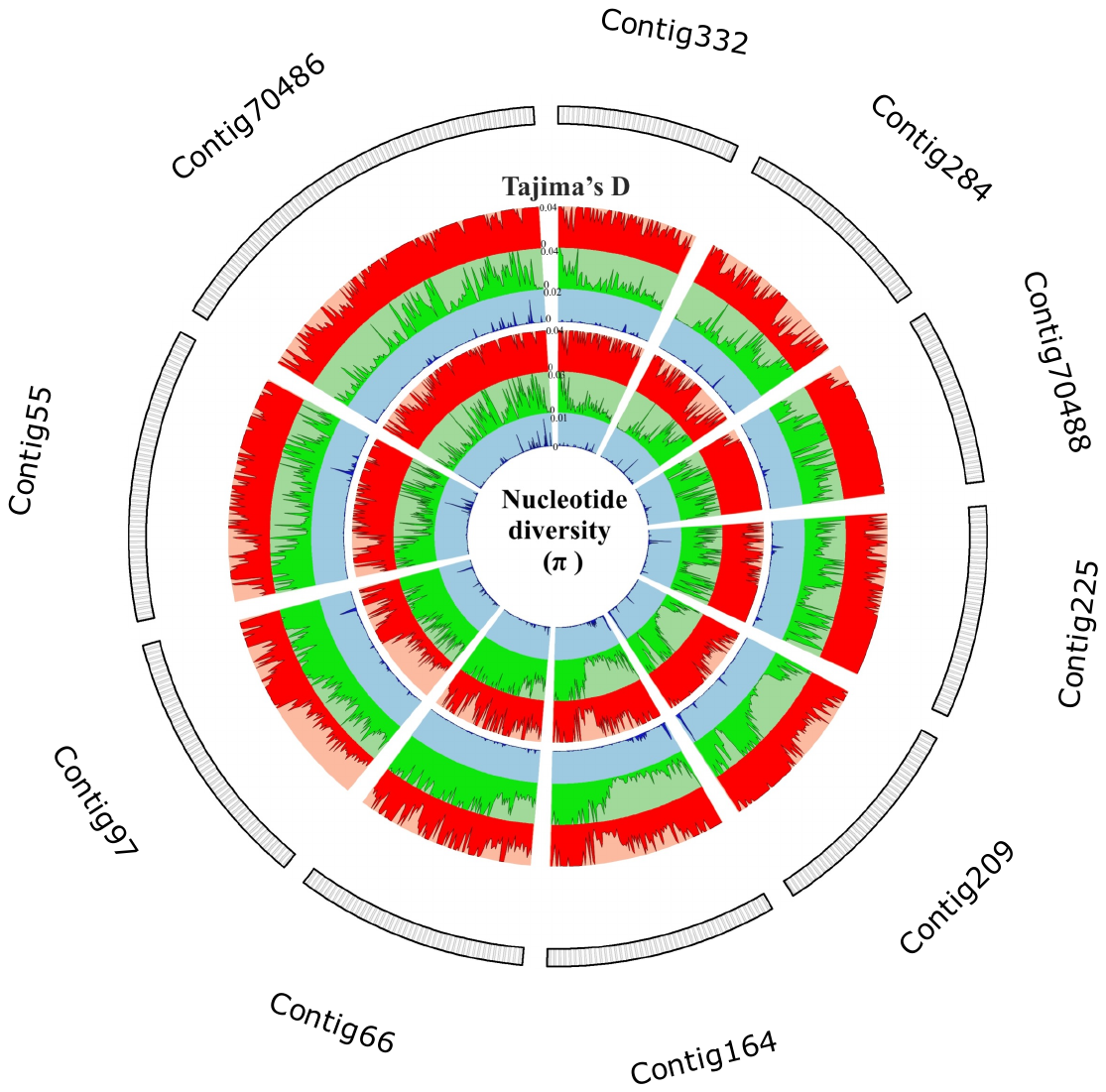

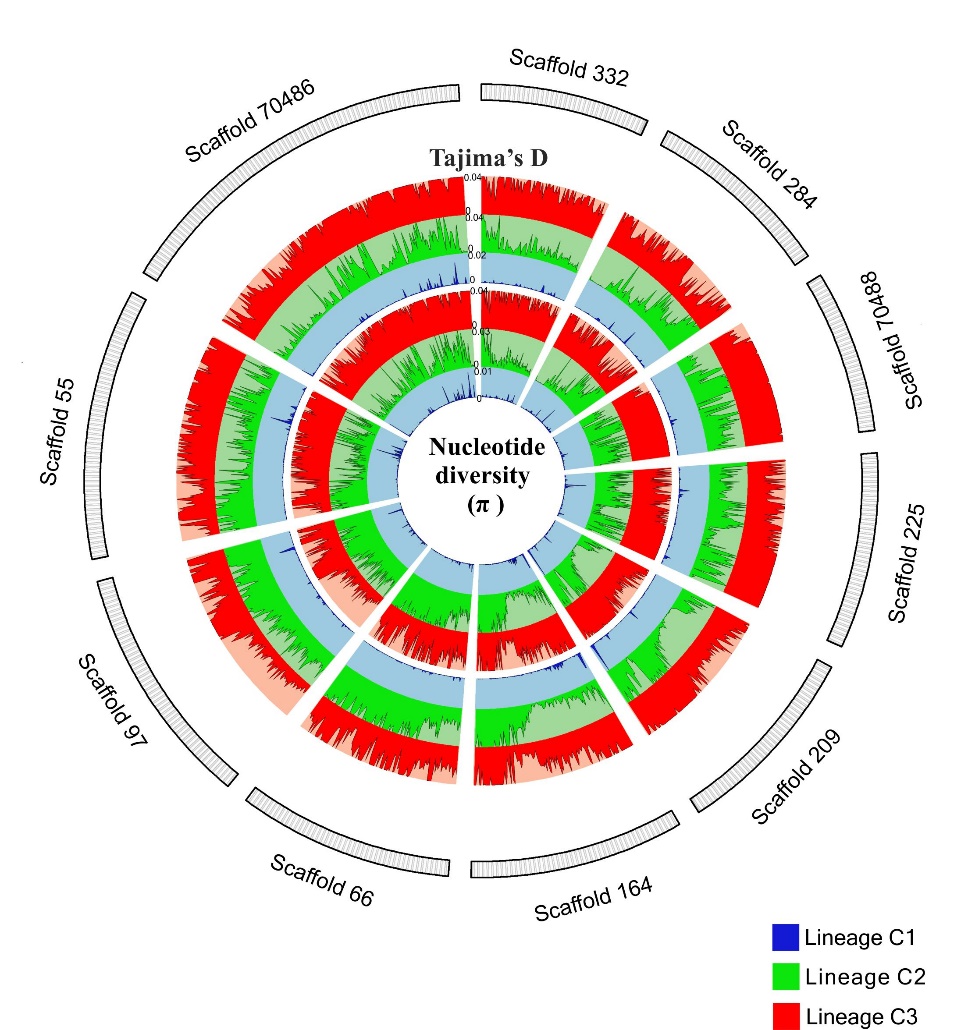
**Figure S3.** Circos plot showing genome-wide diversity of the ten largest contigs of *Colletotrichum truncatum* genomes. Nucleotide diversity (π) per SNP and Tajima’s D were calculated in sliding windows of 10 kb.
