## Supplementary material for "Means, motive, and opportunity for biological invasions: genetic introgression in a fungal pathogen": Figure S4

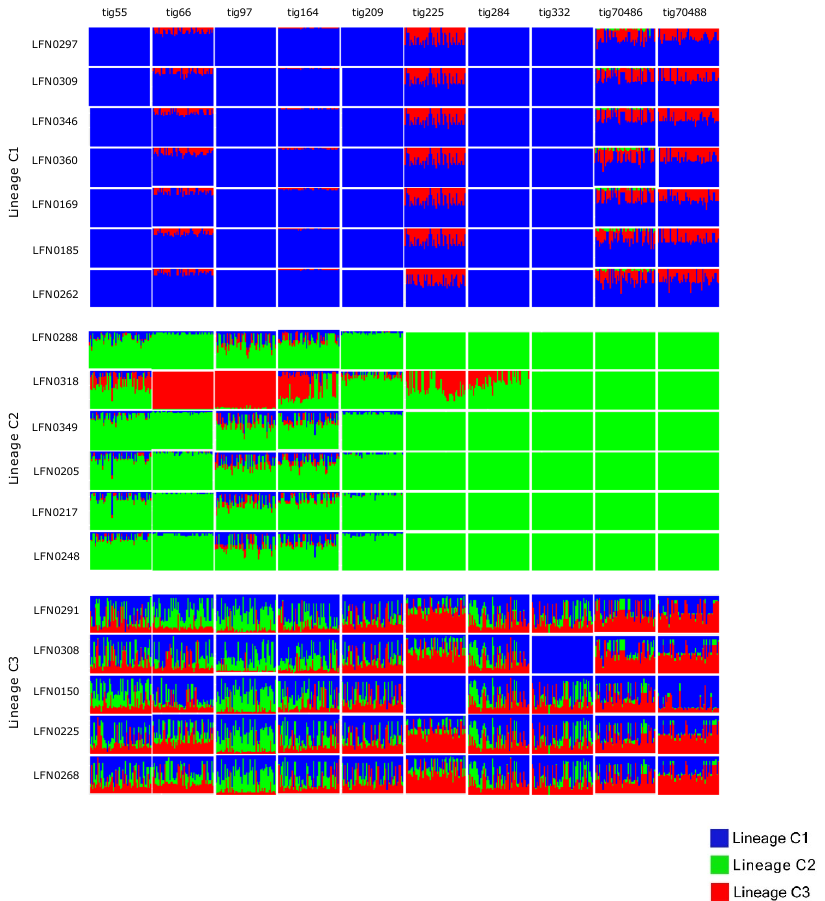

**Figure S4.** Probabilistic chromosome painting bar plots of the ten largest contigs among three *Colletotrichum truncatum* lineages (C1, C2, and C3).
