## Supplementary material for "Means, motive, and opportunity for biological invasions: genetic introgression in a fungal pathogen": Figure S5

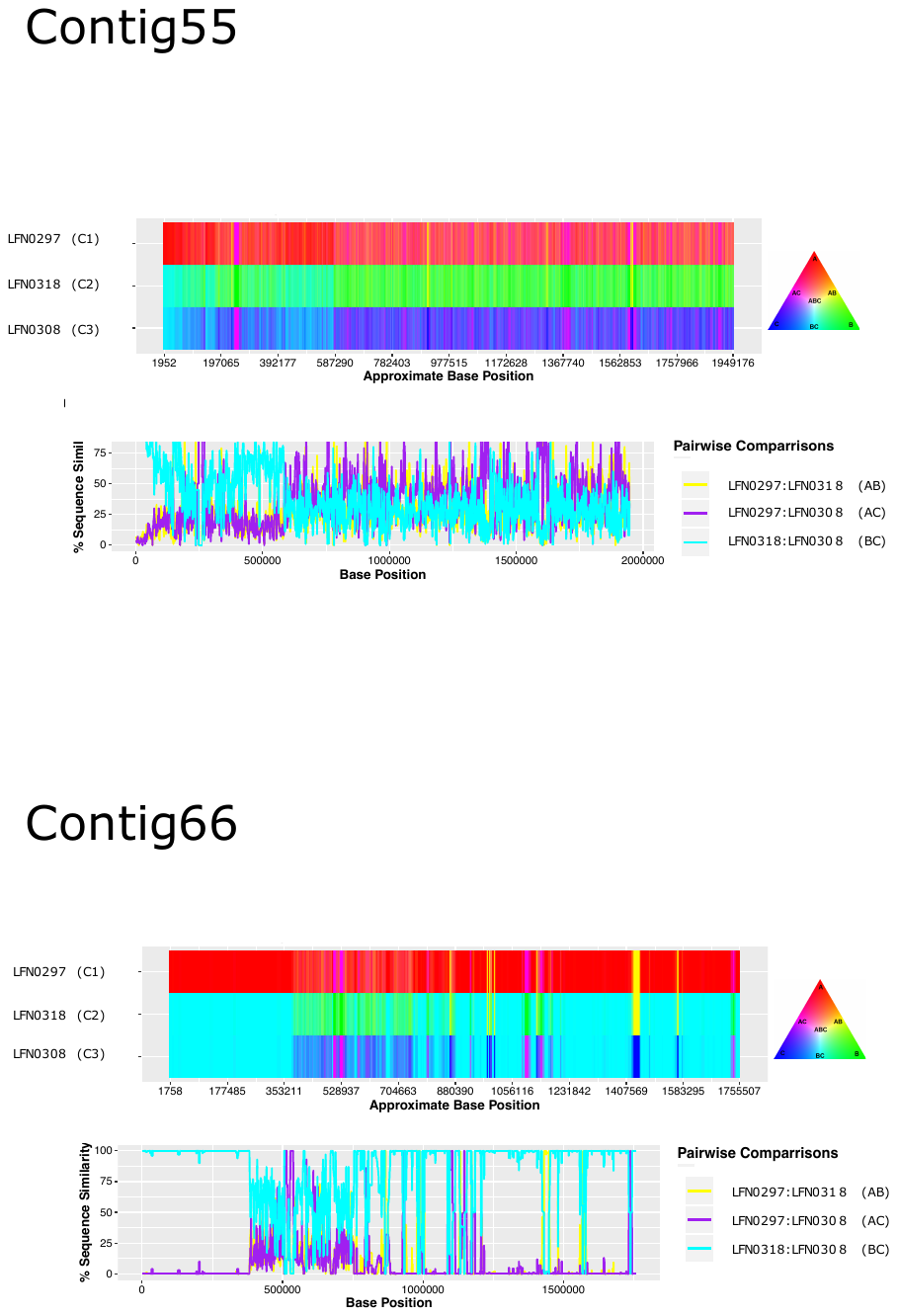

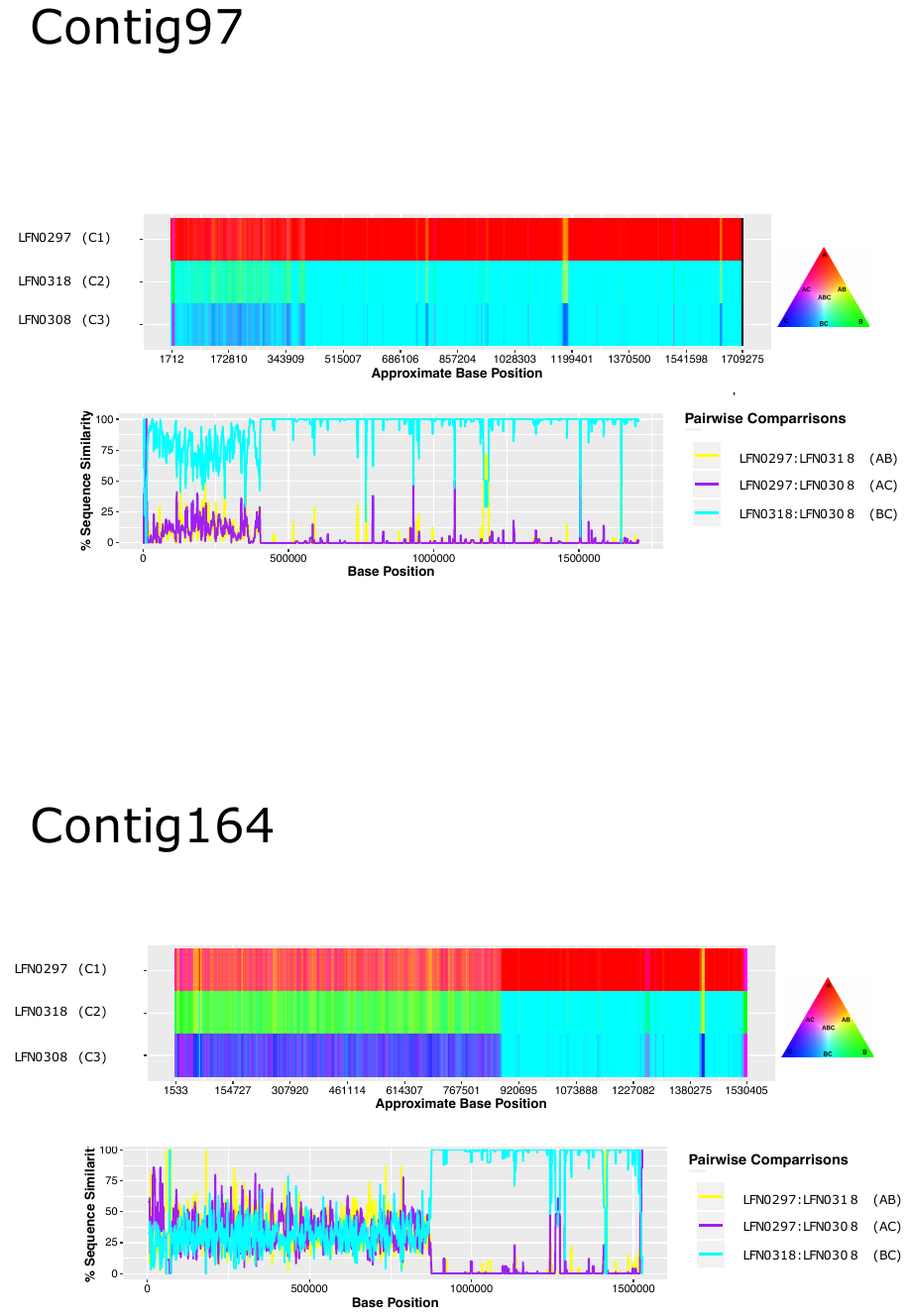

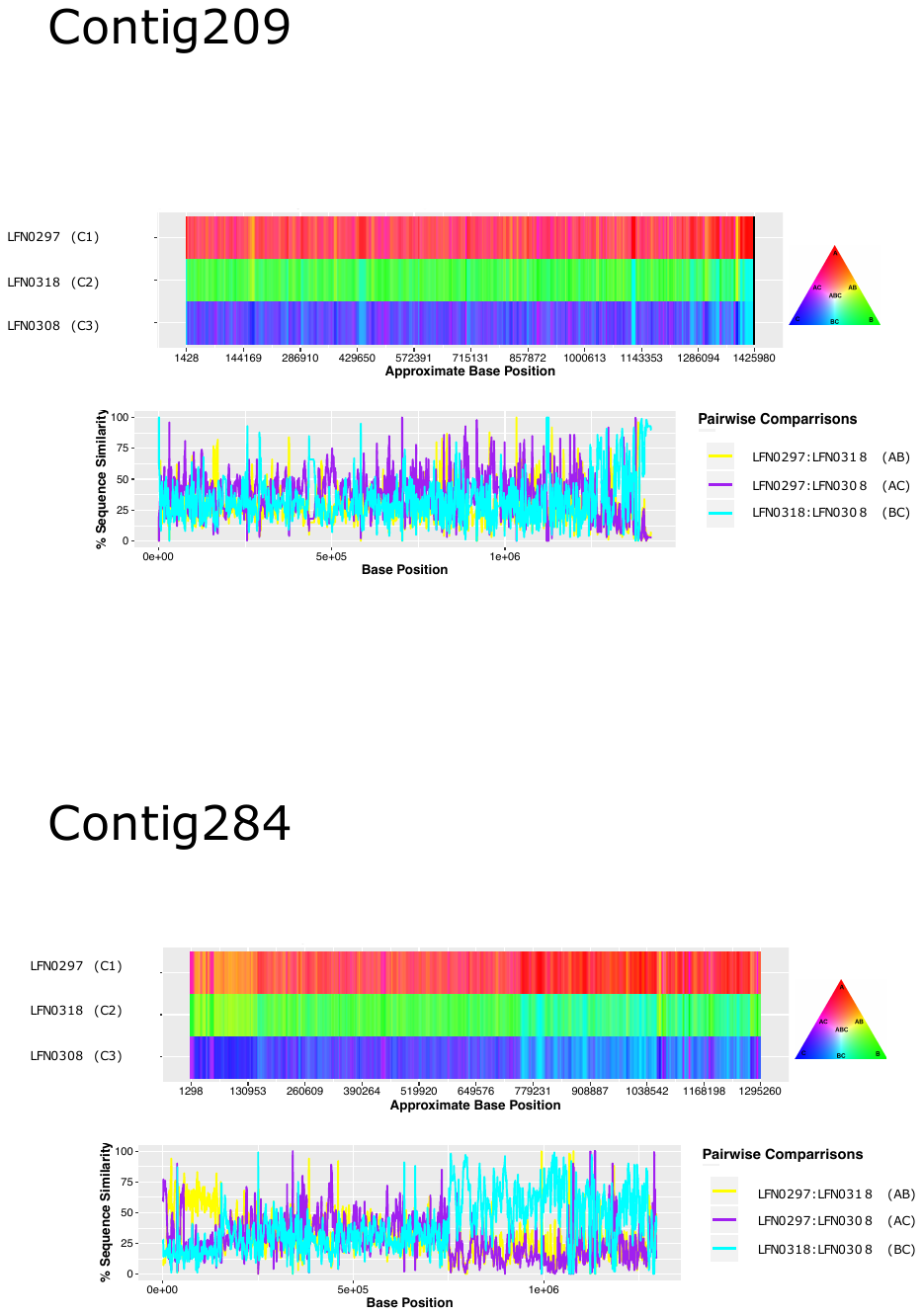

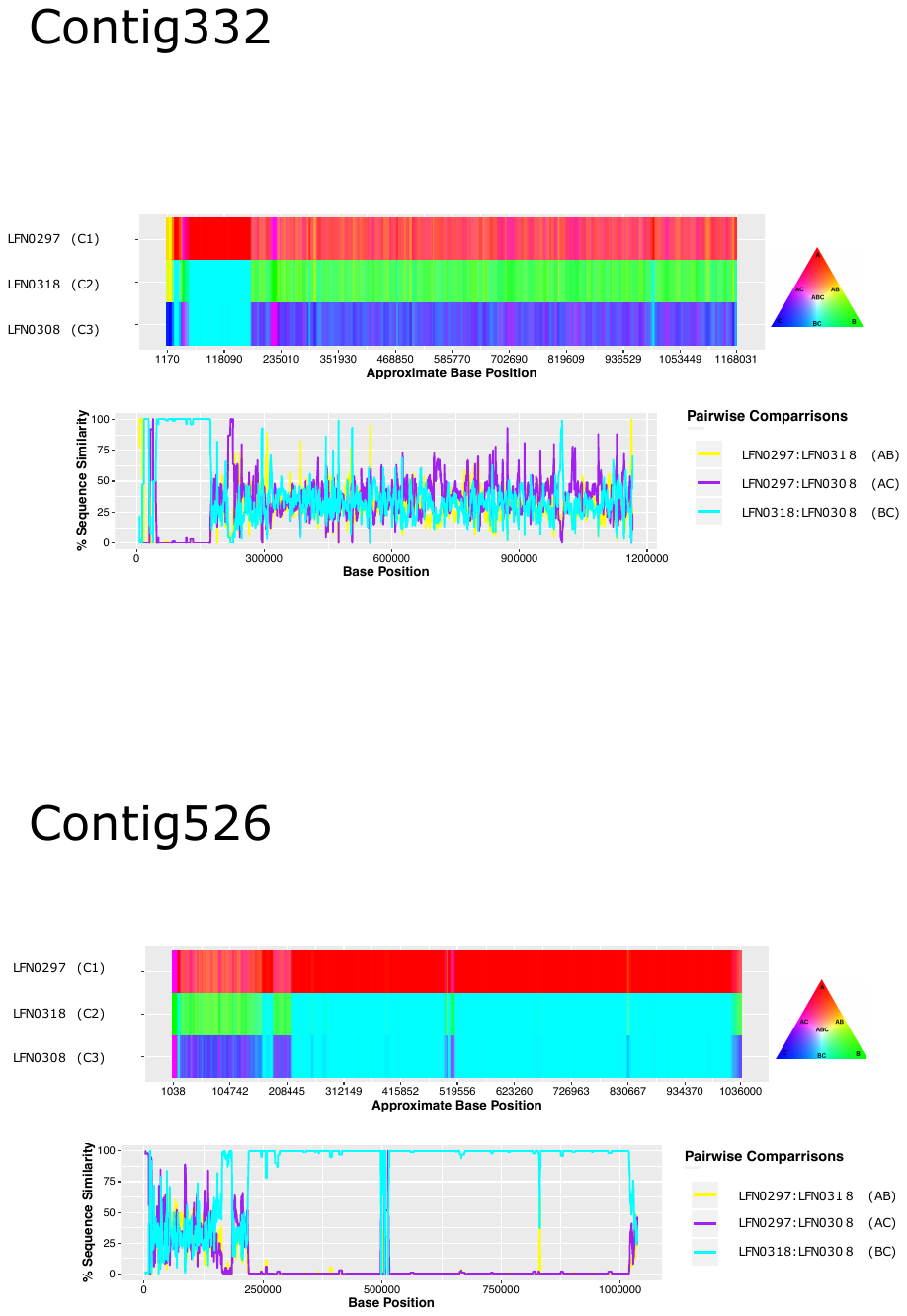

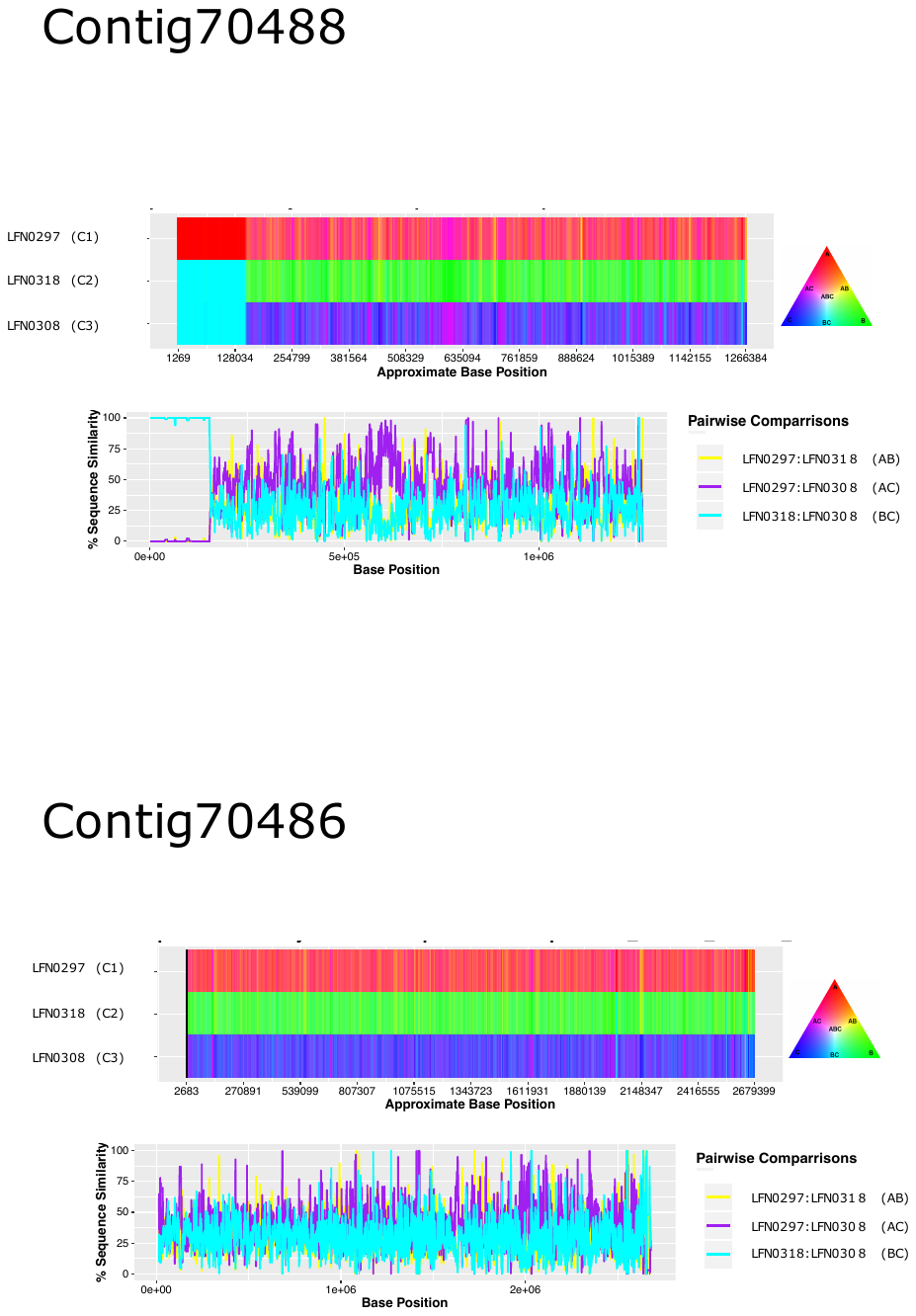

**Figure S5.** Variation in sequence similarity between *Colletotrichum truncatum* lineages (C1, C2, and C3) on the ten largest contigs. The isolates LFN0297 (lineage C1), LFN0318 (lineage C2), and LFN0308 (lineage C3) were used as representative of each lineage and visualized as mosaic-like genome structure through RBG color triangular in the software HybridCheck.
