## Supplementary material for "Means, motive, and opportunity for biological invasions: genetic introgression in a fungal pathogen": Figure S6

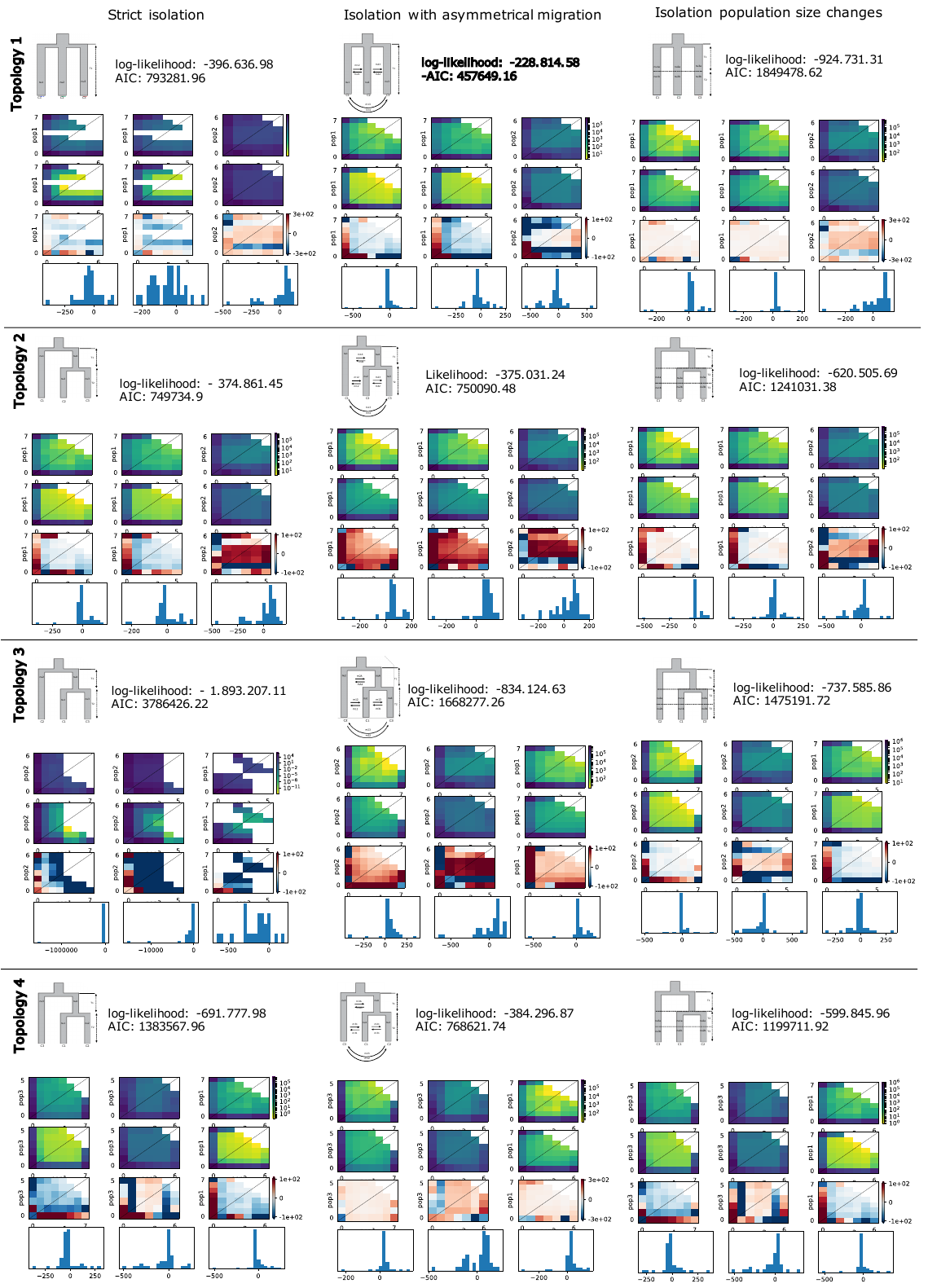

**Figure S6.** Demographic models tested using the three-dimensional site frequency spectrum (3D-SFS) between *Colletotrichum truncatum* lineages (C1, C2, and C3). Data, models and residuals are presented using a heatmap. The best model fitted is in bold. The following parameters were estimated: nu1: size of C1 after split; nu2: size of C2 after split; nu3: size of C3 after split; m12: asymmetrical migration between C1 and C2; m21: asymmetrical migration between C2 and C1; m23: asymmetrical migration between C2 and C3; m32: asymmetrical migration between C3 and C2.; m13: asymmetrical migration between C1 and C3; m31: asymmetrical migration between C3 and C1; T1: scaled time between the split and the size change (in units of 2*Na generations). LL = log likelihood of the model. AIC: Akaike information criterion.
