## Supplementary material for "Means, motive, and opportunity for biological invasions: genetic introgression in a fungal pathogen": Table S1

**Table S1.** Strains and genome sequencing information

| **Lineage** | **Isolate** | **SRA accession number** | **Coverage** | **Mean depth** |
| --- | --- | --- | --- | --- |
| C1 | LFN0169 | SRX7095339 | 100.0 | 112.0 |
| C1 | LFN0185 | SRX7095348 | 99.9 | 115.0 |
| C1 | LFN0262 | SRX7095353 | 100.0 | 150.5 |
| C1 | LFN0309 | SRX7095343 | 99.9 | 56.7 |
| C1 | LFN0360 | SRX7095347 | 100.0 | 154.6 |
| C1 | LFN0297 | SRX7095341 | 100.0 | 137.4 |
| C2 | LFN0346 | SRX7095345 | 87.7 | 104.4 |
| C2 | LFN0205 | SRX7095349 | 83.7 | 76.5 |
| C2 | LFN0217 | SRX7095350 | 86.0 | 109.3 |
| C2 | LFN0248 | SRX7095352 | 84.6 | 77.0 |
| C2 | LFN0318 | SRX7095344 | 99.9 | 102.5 |
| C2 | LFN0349 | SRX7095346 | 86.1 | 106.9 |
| C3 | LFN0288 | SRX7095355 | 84.8 | 103.7 |
| C3 | LFN0150 | SRX7095338 | 83.9 | 83.1 |
| C3 | LFN0225 | SRX7095351 | 79.8 | 97.9 |
| C3 | LFN0268 | SRX7095354 | 87.5 | 89.0 |
| C3 | LFN0291 | SRX7095340 | 86.2 | 109.9 |
| C3 | LFN0308 | SRX7095342 | 84.9 | 98.6 |
